# An inverse correlation between linguistic and genetic diversity

**DOI:** 10.1101/2024.12.18.628602

**Authors:** Anna Graff, Erik J. Ringen, Taras Zakharko, Mark Stoneking, Kentaro K. Shimizu, Balthasar Bickel, Chiara Barbieri

## Abstract

Human population history, as traced by our genome, has shaped the distribution of languages around the world. While case studies suggest that population history can also affect specific structures of languages, such as patterns in their sound systems and grammars, results are conflicting, and it remains unknown whether such effects hold globally. Here we show that, adjusting for geography, phylogeny, and environment, genetic diversity is inversely correlated with diversity in linguistic structures. Low genetic diversity (i.e. excess homozygosity) results from relative isolation, and this promotes diversification in language. High genetic diversity results from contact and migration, and this promotes homogenization in language. These effects are particularly pronounced in Asia and, under some conditions, also in a few other areas, suggesting sociocultural factors contributing to differences in how strongly isolation favors linguistic diversity and how strongly contact promotes linguistic homogenization. Our results suggest that present-day hotspots of linguistic diversity, which are characterized by relative isolation, may provide a privileged window on the dynamics of linguistic evolution as they are less affected by the vast demographic expansions, migrations, and extinctions that characterize the Neolithic age.

## Main Text

The genetic patterns that emerge from human population history explain much of the distribution of languages over space and time, although not to the extent that Darwin surmised in his famous proposal that the human pedigree would match the phylogeny of languages (*1*). Much less is known about how genetic patterns also explain the distribution of the structures that languages have, such as patterns in their grammars, vocabularies, and sound systems, which tend to be similar over larger arrays of languages (Fig. 1). Genetic distances are correlated with grammar in Northeast Asia (*2*), vocabulary in Central African hunter-gatherers (*3*), and sound systems at a global scale (*4*). However, it remains an open question how general such correlations are across different language structures and world regions. While correlations with patterns of sound (*5, 6*) and meaning (*7*) might in rare cases be partially driven by functional relationships with genes, it is unknown whether other mechanisms beyond shared local history impact language diversity. One candidate mechanism is grounded in the dynamics of isolation and contact.

**Fig 1.**
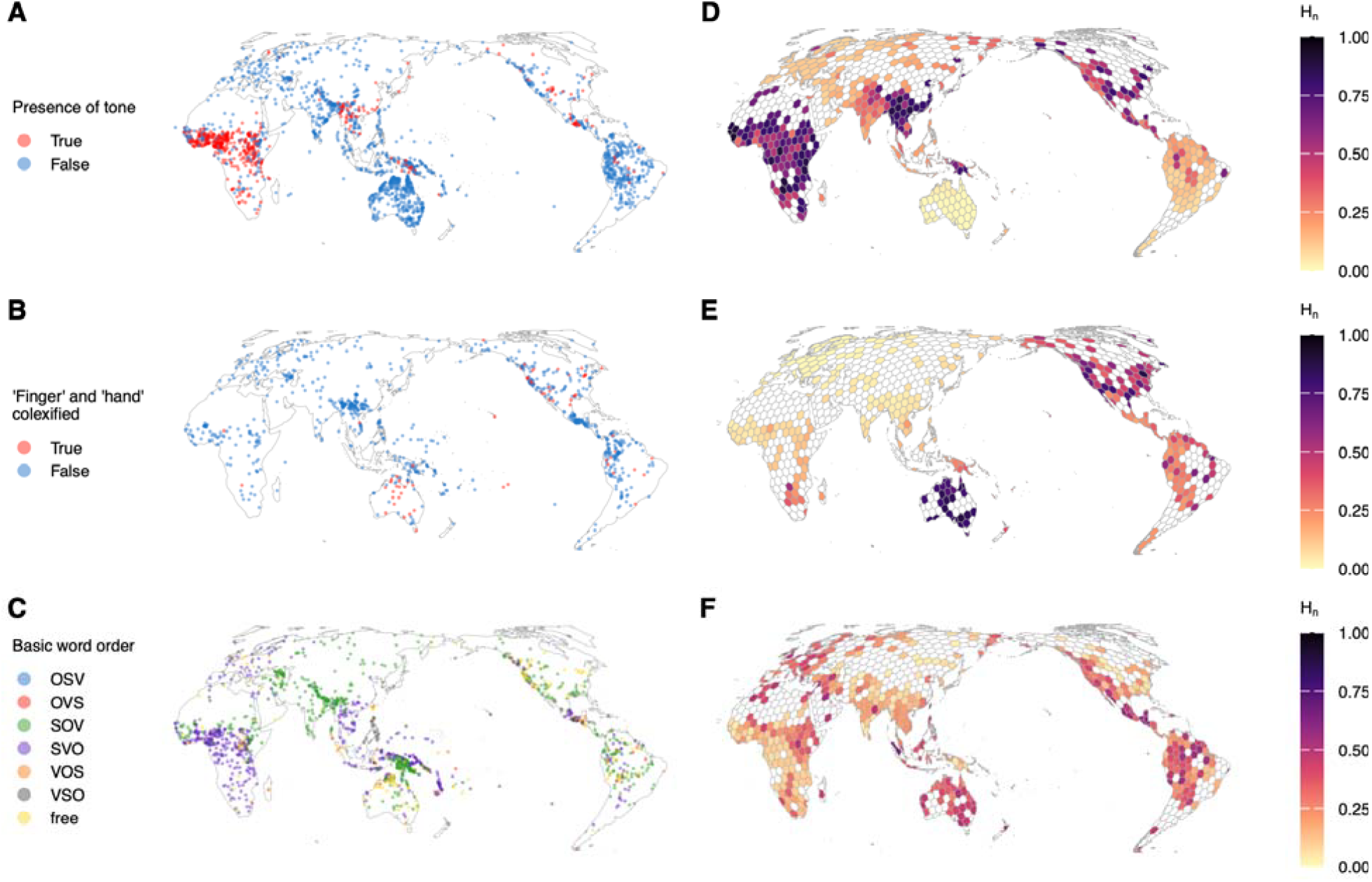
Examples of the uneven distribution of global structural diversity for three linguistic features from different linguistic domains. The left-hand maps show each coded language’s feature value. The right-hand maps show the model-based estimates of normalized entropies (H_n_) of each feature’s distribution, adjusted for spatial and phylogenetic autocorrelation. Grid cell diameters measure 500 km. (**A–B**) Presence of tone (phonology) in N=2236 languages. (**C–D**) Use of the same word (“colexification”) for “finger” and “hand” (lexical semantics) in N=881 languages. (**E–F**) Basic word order (S: most agent-like noun phrase, O: most patient-like noun phrase, V = verb; morphosyntax) in N=1502 languages.

Human populations over the millennia have oscillated between contact and isolation, impacted by bouts of demic dispersals and expansions which left discernible traces in the genome (*8*). It has been hypothesized that contact and isolation can directly impact linguistic structures, with contact leading to homogenization through horizontal borrowing and isolation favoring diversification through vertical evolution (*9*). Specifically, high levels of migration and wide-spread population contact are expected to result in regions of reduced diversity in structures (*spread zones)*. In contrast, structurally diverse regions (*accumulation zones*) would emerge in areas where linguistic diversity is maintained or enriched, often in contexts of limited integration and increased isolation at the fringes of spread zones (*9*–*11*). From this perspective we expect linguistic homogenization to correlate with more genetic diversity (at the individual level) in populations under intensified contact, and linguistic diversification to correlate with less genetic diversity, caused by isolation under reduced contact. It is currently unknown whether and to what extent this correlation holds at a global scale.

We assess the correlation between genetic diversity and a wide range of linguistic structural diversity by combining genetic data from 5770 individuals in 650 populations with linguistic data from 4255 languages sampled worldwide, coded for 333 structural features. Several confounds need to be adjusted for because the distribution of languages, and therefore their structures, is also heavily driven by the natural environment, population sizes and densities as well as by socio-cultural practices, which have shaped pathways of linguistic diversification (Fig. 2) (*12*–*22*). Most of these effects reflect shared history although the relationship between sound systems and environmental temperature (*23*–*25*) and subsistence type (*26*) are possibly adaptive. Further confounds are phylogenetic and spatial autocorrelation (*21, 27*).

**Fig 2.**
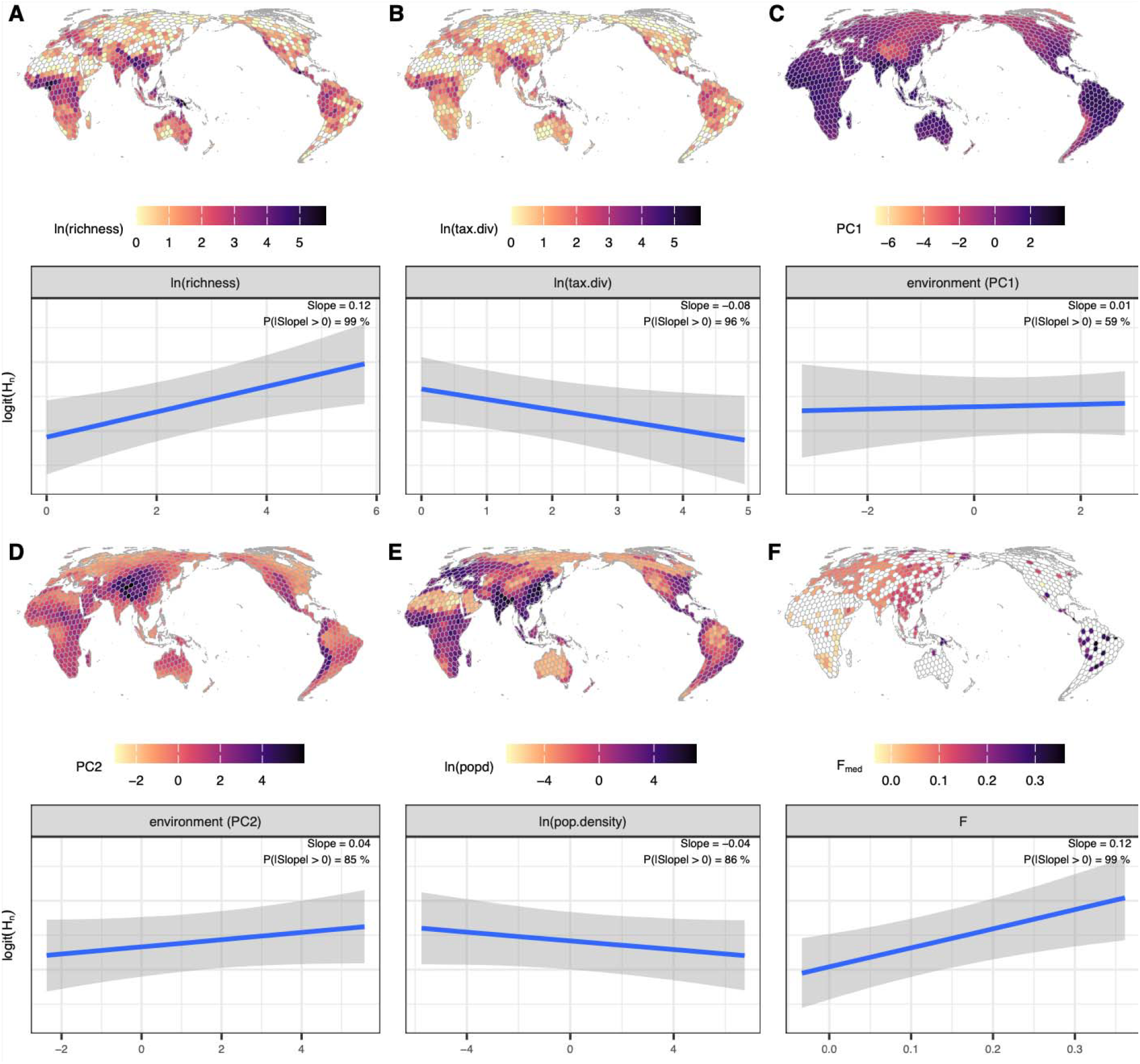
Distributions and marginal effects of the six main predictors on cell-wise linguistic diversity (H_n_, normalized entropy) when the others are at their mean in the best-fitting model. Intervals correspond to the 89% credible intervals. (**A**) (Log) language richness, i.e. number of languages and dialects per cell. (**B**) (Log) taxonomic diversity of languages and dialects per cell, scoring how diverse and balanced the included languages’ taxonomic relationships are. (**C**) First PC of environmental predictors (R^2^ = 34.8). Main loadings are the number of months with mean temperature > 15°C (27.3%), mean annual temperature (23.7%) and temperature of the warmest quarter (22.7%). (**D**) Second PC of environmental predictors (R^2^ = 18.6). Main loadings are altitude (26.6%), altitude variation (20.1%) and seasonal variance of precipitation (15.6%). (**E**) (Log) Population density. (**F**) Median Wright’s *F* coefficient, with increasing values indicating higher excess homozygosity and therefore lower levels of genetic diversity.

### Modelling diversity estimates for linguistic features

We probe the extent to which linguistic diversity is driven by genetic and ecological predictors across the cells of a geodesic grid that partitions the world’s land mass into near-uniformly spaced hexagons (*28, 29*). A median cell diameter of 500 km reduces the uncertainty of exact language locations and captures the potential for demographic, cultural, and linguistic contact, whilst ensuring adequate data coverage per cell. As a sensitivity analysis, we also consider a smaller grid with a median cell diameter of 300 km (fig. S1).

We estimate linguistic diversity in terms of normalized Shannon entropy for 333 features relating to phonology, lexical semantics and grammar (see Fig. 1 for one example each) from a dataset that is curated for logical and statistical independence between features (Typology Linked and Independent “TLI-statistical”) (*30*). The cell-wise entropy estimates are obtained though Bayesian generalized additive mixed models (GAMMs) that adjust for universal baseline expectations across all cells, the geohistorical area of each cell (*31*) and phylogenetic and spatial autocorrelations between the cells (*Supplementary Materials*).

Model-based entropy estimates are then used as the (logit-transformed) response variable for a set of eight GAMMs (*Supplementary Materials*; fig. S2 for posterior predictive checks on all models), designed to quantify the marginal effect of genetic diversity on cell-wise structural linguistic diversity, adjusting for the following predictors: (log) language richness, i.e. number of languages; (log) linguistic taxonomic diversity, scoring how diverse and balanced the included languages’ taxonomic relationships are; environmental variation including climatic, altimetric, productivity and land use factors; (log) population density from historical estimates; geohistorical areas delineated in the AUTOTYP database (*31*); and genetic diversity, represented by excess homozygosity per individual, i.e. the *F* coefficient, where high values of *F* correspond to low genetic diversity (table S1 and Fig. 2). We further include adjustments over the geographic location of each grid cell to control for spatial autocorrelation. Our null model m1 considers only language richness and taxonomic diversity as predictors (Fig. 2a–b). Models m2-m4 further include the first two principal components of environmental predictors, with PC1 mainly loaded by variables related to climate, and PC2 mainly loaded by variables related to terrain (m2; Fig. 2c–d); human population density (m3; Fig. 2e) or the degree of genetic diversity sampled within each grid cell, expressed by the median of the individual coefficient *F* (m4; Fig. 2f). Models m5-m7 further include additive pair-wise combinations of these predictors, and the full model m8 includes all of them. By including random slopes by geohistorical area in each model, we both allow for socio-culturally conditioned variations in the impact of the predictors and account for deep population history with regard to the genetic predictor (which globally displays the characteristic out-of-Africa gradient of *F*; Fig. 2f). Feature-specific effects are also accounted for throughout our models via random intercepts and slopes. Approximate leave-one-out cross-validation (*32*) shows that across conditions, the full model m8 including all predictors strongly outperforms all other models (all Δ_elpd_ > 136 ± 13, see fig. S3).

### Genetic and geographical isolation predict high structural diversity in language

High *F* values, i.e. low median individual-level genetic diversity, are the strongest and most unambiguous predictor of high structural linguistic diversity (posterior probability of a positive effect is 99%; Fig. 2f). This means that an increase in the (scaled) *F*-predictor by one standard deviation is associated with a 0.12 increase in the logit-transformed normalized Shannon entropy (denoting structural linguistic diversity), roughly corresponding to an increase in normalized Shannon entropy by 2.5%, when all other predictors are held at their mean. Low genetic diversity is the result of small effective population size, isolation and lack of gene-flow between groups. Our results show that these demographic factors promote diversification and/or impede homogenization in language, consistent with the hypothesized correlation. The result is robust at a higher spatial resolution (smaller grid cells), and it is also apparent when using an alternative linguistic dataset (the Grambank Independent, “GBI-statistical” dataset, see Methods), albeit with less posterior certainty because of reduced data coverage (fig. S4). The effect is similar in all ten geohistorical areas (with all posterior probabilities of a positive effect Δ91%; Fig. 3 and fig. S5), but it is particularly strong in Asia and the New Guinea – Oceania area, a pattern that is replicated in the smaller-grid analysis (fig. S6) but has more limited scope in the GBI dataset (figs. S5-6) and also includes South Central America in the main, but not other analyses. A possible explanation of this increase might be active processes of cultural compartmentalization and linguistic divergence (*33,9,10,34*–*40*), but denser global data coverage is needed to test this possibility against other demographic and social processes.

**Fig 3.**
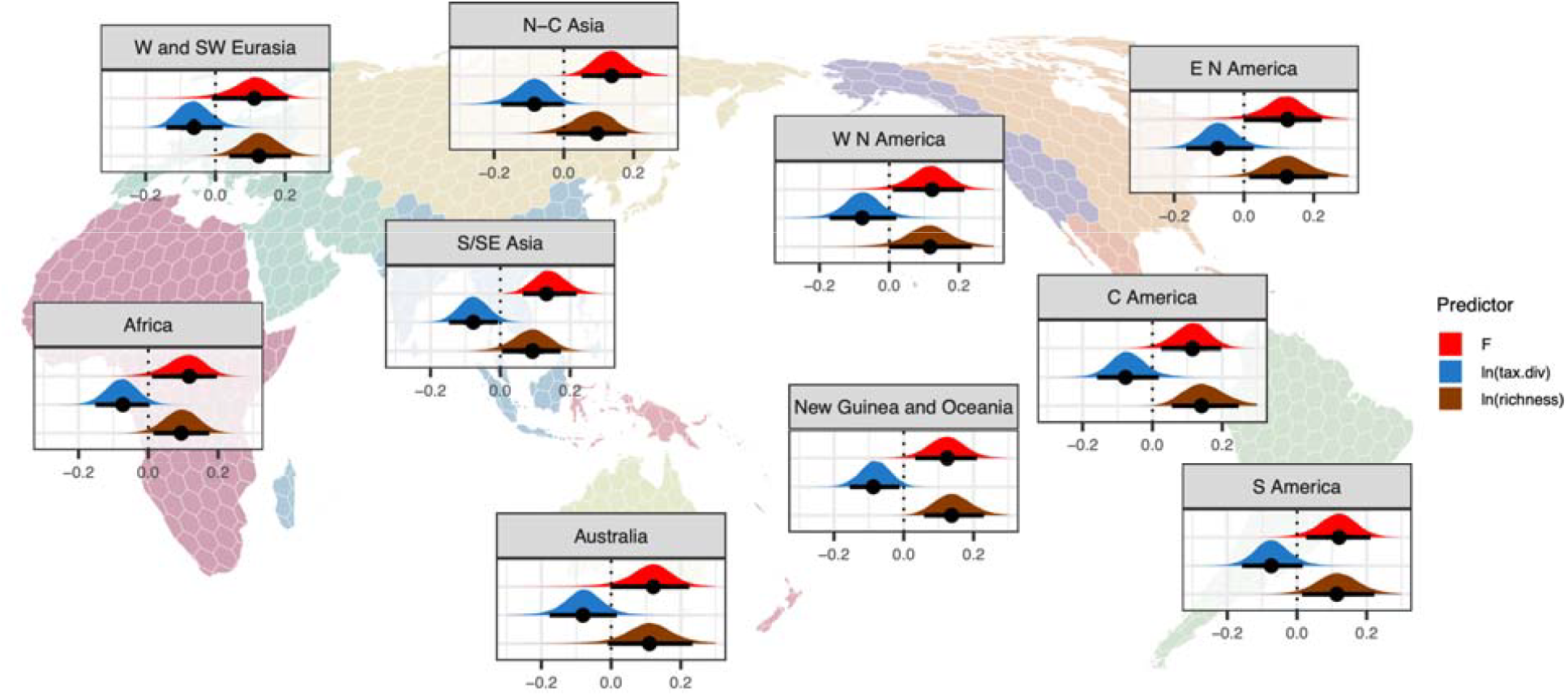
Group-level marginal effects (slopes) of the effects with strongest size and support (genetic diversity *F*, log taxonomic diversity and log language richness). by geohistorical area as identified by AUTOTYP (*31*). The interval bars correspond to the 89%-HPDI.

Aside from low genetic diversity, language richness but not taxonomic diversity enhances structural diversity (marginal effects at the mean of 0.12 for richness, roughly an increase in normalized Shannon entropy by 2.5%, with a 99% posterior probability of a positive effect; -0.08 for taxonomic diversity, roughly a decrease in normalized Shannon entropy by 1.6%, with a 96% posterior probability of a negative effect; Fig. 2a–b). This result is also particularly robust in certain continents (fig. S5) in the main analysis. Although our sensitivity analyses (fig. S4–6) do not replicate this finding, it tentatively suggests that high language density favors structural diversity, but more so when languages are related than in the presence of high taxonomic diversity. This is consistent with observations of divergence when related languages are in contact (*35,41,39,*), an effect known as schismogenesis (*42, 43*). Structural diversification may be more relevant among speakers of related languages since they share common ethnolinguistic histories, and may therefore be at stronger need to mark social differences (*44*) than more distantly related neighbors.

Past studies highlighted that group size, population density, climate and mountainous terrain each have a relevant but differential impact on language richness (*16*–*18,21,20*). While our results generally favor the notion that higher population densities – associated with more local contact – decrease diversity in language structure (marginal effect at the mean: -0.04, roughly a decrease in normalized Shannon entropy by 0.9%), the effect has only 86% posterior probability (Fig. 2e) and it is not consistent across all sensitivity analyses (fig. S4). Similarly, regarding the environmental variables loading PC1, our results provide no support (marginal effect at the mean: 0.01, roughly a decrease in normalized Shannon entropy by 0.3%; 59% posterior probability of a positive effect) for factors associated with warm climates to favor structural diversification (Fig. 2c). For increases in environmental PC2 – mainly loaded by variables indicative of mountainous terrain, which has sometimes been proposed to impede cultural contact and to harbor linguistically “conservative” and highly diverse regions (*45*) – we find increases of structural diversity (marginal effect at the mean of 0.04, roughly an increase in normalized Shannon entropy by 0.9%). However, the posterior probability of a positive effect is only 85% (Fig. 2d). More extensive datasets – particularly with respect to genetics – could resolve the currently inconclusive effect directions, sizes and support of these predictors. The fact that models excluding the environmental PCs and population density as predictors yield less predictive accuracy than the full model suggest that they cannot be dismissed as drivers of structural diversity.

### Effects on linguistic diversity vary by feature

The effects of genetic diversity *F*, language richness and taxonomic diversity, the three effects with strongest size and support, vary substantially across features (Fig. 4). 23% of features have positive 89%-HPDIs that exclude zero for *F* (9% of features have negative effects and 68% of features include zero in their 89%-HPDI). For language richness, these percentages lie at 18%, 5% and 77%, respectively, and for taxonomic diversity they amount to 6%, 19% and 75% (Fig. 4c). Further research is needed to probe potential factors that explain this variation, but we note that there is no apparent correlation with domains of language (Fig 4a, figs. S7a, S8a). Notably, our findings also do not support a correlation with features of language morphology (word forms) (Fig. 4b, figs. S7b, S8b) that have been proposed to homogenize towards simplification in the presence of second-language learners, i.e. intensified contact (*46*–*50,4,51,52*). We also found no patterns of such features with population density, an alternative proxy for contact (figs. S7b, S8b) (*53*). This is consistent with other recent challenges to the notion that contact leads to homogenous simplification (*54*–*58*).

**Fig 4.**
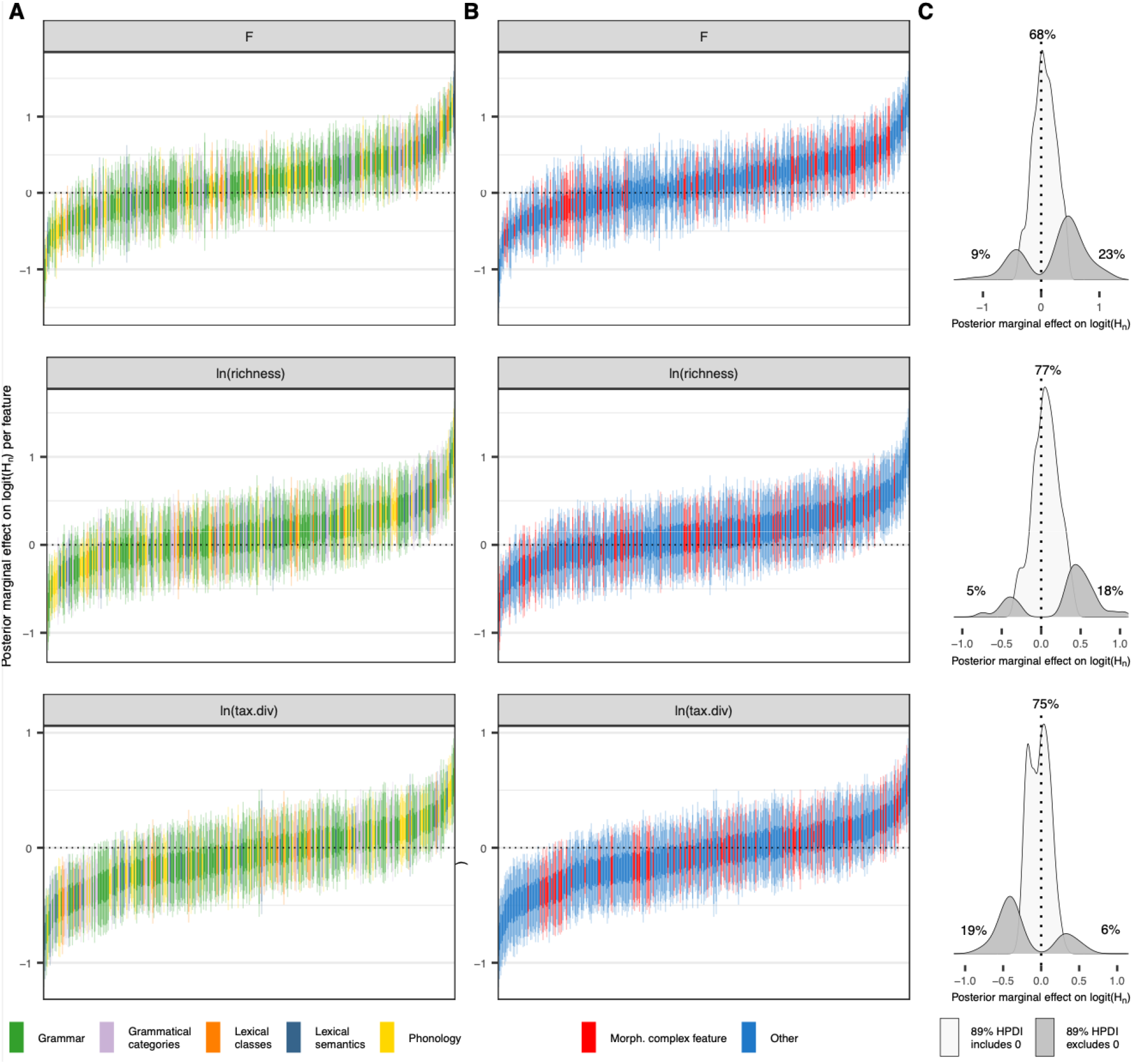
Group-level marginal effects (slopes) of the effects with strongest size and support (*F*, log language richness and log taxonomic diversity) by feature. (**A**) Highest posterior density intervals (HPDIs; 50%, 89% and 95%), ordered by increasing marginal effect for each predictor and colored by feature domain (grammar, grammatical categories, lexical classes, lexical semantics and phonology; as provided in the TLI dataset) (*30*). (**B**) HPDIs (50%, 89% and 95%), ordered by increasing marginal effect for each predictor, with features pertaining to morphological complexity (in particular, synthesis and the degree of fusion and/or affixation) (*30 48 56*) indicated by red coloring. The list of features can be found in the OSF repository. Neither feature domain nor morphological complexity are associated with differential effects on linguistic diversity. (**C**) Density plots of features with the 89% HPDI excluding vs. including zero for each predictor.

### Structural diversity hotspots as potential gateways to past linguistic landscapes

The tight link between human population history and structural linguistic diversity has implications for our understanding of linguistic diversity in space and time. In our data, contact-ridden regions – those associated with high genetic diversity – exhibit low levels of linguistic diversity. This is consistent with the hypothesis of spread zones, which would mostly have emerged as a consequence of demic and/or cultural spreads of language families associated with Neolithic transitions to food production and, more recently, to imperial or colonial states. In contrast, in our data, present-day hotspots of linguistic diversity are associated with low genetic diversity (excess homozygosity) and, therefore, demographic histories with relatively limited and/or different patterns of contact, supporting the notion of accumulation zones (*9*–*11*). Languages in present-day diversity hotspots would of course have evolutionary histories shaped by descent, innovation and contact effects like any other language, but they would have experienced less massive turnovers through extensive contact in comparison to spread zones. As a result, they may not only better reveal the breadth of structural linguistic variation than spread zones but they could also shed light on the dynamics of structural linguistic evolution before the Neolithic turnovers, when population densities were lower and contact less driven by mass expansions and migrations. Thus, to exclude the confounding effects of recent historical dynamics, linguistic research at the level of the human species needs to focus more actively on diversity hotspots. This move is urgent in the face of the rapid global decline of language diversity (*59, 60*).

## Acknowledgments

We thank Chundra Cathcart for discussions on diversity hotspots.

## Author contributions

Conceptualization: AG, MS, KKS, CB, BB. Data curation: AG, TZ, CB, BB. Methodology: AG, ER, BB. Investigation: AG. Visualization: AG, ER, BB. Validation: AG, ER. Funding acquisition: KKS, BB. Project administration: AG. Supervision: KS, CB, BB. Writing – original draft: AG. Writing – review & editing: AG, ER, TZ, MS, KKS, CB, BB.

## Data and materials availability

All models and analyses presented here as well as the data, files and scripts necessary to reproduce them are available at OSF (https://osf.io/2qgje/).

