## Supplementary Materials for "An inverse correlation between linguistic and genetic diversity"

**Supplementary Materials for:
An inverse correlation between linguistic and genetic diversity**

Anna Graff, Erik J. Ringen, Taras Zakharko, Mark Stoneking, Kentaro K. Shimizu, Balthasar Bickel, Chiara Barbieri

Materials and Methods

Features, grids and diversity estimates

Linguistic features were drawn from two datasets: TLI-statistical (333 features in 4257 languages overall) and, for a sensitivity analysis, GBI-statistical (196 features in 2467 languages overall) (*30*). These datasets were curated to reduce logical and statistical dependencies between features within each dataset. For each feature, all available languages were modelled in a GAMM implemented using the brms (*61*) interface to Stan (*62*, *63*).

We used the approach formalized by Rivière (*28*) and implemented by Zakharko (*29*) to create a geodesic hexagonal grid used for assigning languages to geographic units via their Glottolog point coordinates in all models in R (*64*). For the main analysis, the generated grid measured a median cell diameter of 500 km, producing 1140 cells across the world’s land mass (including islands). An alternative grid with median cell diameter of 300 km (2833 cells) was produced for the sensitivity analysis.

The TLI and GBI datasets each include a majority of binary features and a minority of categorical, multistate features. These were modeled differently. The probability of each binary feature (N=269 in TLI, N=193 in GBI) was modelled for every language *i* as follows:

$$\mathrm{state}_{i}\sim Bernoulli\left( \pi_{i} \right)$$

$$\pi_{i}=\frac{exp\left( \eta_{i} \right)}{1+exp\left( \eta_{i} \right)}$$

$$\eta_{i}=\alpha+\alpha_{\mathrm{AREA}_{i}}+\alpha_{GRID.\mathrm{ID}_{i}}+\alpha_{\mathrm{FAMILY}_{i}}+t2(\mathrm{lon}_{GRID.\mathrm{ID}_{i}},\mathrm{lat}_{GRID.\mathrm{ID}_{i}})$$

$$\alpha\sim N\left( 0, 2 \right)$$

$$\alpha_{\mathrm{AREA}_{i}}\sim N\left( 0, \sigma_{\mathrm{AREA}_{i}} \right)$$

$$\alpha_{GRID.\mathrm{ID}_{i}}\sim N\left( 0, \sigma_{GRID.\mathrm{ID}_{i}} \right)$$

$$\alpha_{\mathrm{FAMILY}_{i}}\sim N\left( 0, \sigma_{\mathrm{FAMILY}_{i}} \right)$$

$$\sigma_{\mathrm{AREA}_{i}}, \sigma_{GRID.\mathrm{ID}_{i},}\sigma_{\mathrm{FAMILY}_{i}}, \sigma_{\mathrm{spline}}\sim N\left( 0, 2 \right)$$

For every language *i,* we estimated the observed feature state. $\alpha$ indicates the intercept, i.e. the global baseline across all grid cells. The terms $\alpha_{\mathrm{AREA}_{i}}$, $\alpha_{GRID.\mathrm{ID}_{i}}$ and $\alpha_{\mathrm{FAMILY}_{i}}$ indicate that the intercept can vary by area, grid cell and family. The term $t2(\mathrm{lon}_{GRID.\mathrm{ID}_{i}},\mathrm{lat}_{GRID.\mathrm{ID}_{i}})$ indicates a tensor product spline for smooths over the geographic location of the centroid of the grid cell the language is located in (coordinates in the Pacific-centered Equal Earth Asia-Pacific projection EPSG:8859). The intercept, all individual varying effects and the standard deviation over the tensor smooth were each assumed to have normal priors with mean 0 and standard deviation  2.

The probability of states in categorical features with J>2 states (N=64 in TLI, N=3 in GBI) in each language *i* were modelled as follows:

$\mathrm{state}_{i}\sim Categorical\left( \pi_{i1}, \pi_{i2}, \ldots, \pi_{iJ} \right)$, where $\sum_{j=1}^{J} \pi_{ij}=1$

$\pi_{ij}$=$\frac{exp\left( \eta_{ij} \right)}{\sum_{j=1}^{J} exp\left( \eta_{ij} \right)}$

$$\eta_{ij}=\alpha+\alpha_{\mathrm{AREA}_{i,j}}+\alpha_{GRID.\mathrm{ID}_{i,j}}+\alpha_{\mathrm{FAMILY}_{i,j}}+{t2}_{j}(\mathrm{lon}_{GRID.\mathrm{ID}_{i}},\mathrm{lat}_{GRID.\mathrm{ID}_{i}})$$

$$\alpha\sim N\left( 0, 2 \right)$$

$$\alpha_{\mathrm{AREA}_{i,j}}\sim N\left( 0, \sigma_{\mathrm{AREA}_{i}} \right)$$

$$\alpha_{GRID.\mathrm{ID}_{i,j}}\sim N\left( 0, \sigma_{GRID.\mathrm{ID}_{i}} \right)$$

$$\alpha_{\mathrm{FAMILY}_{i,j}}\sim N\left( 0, \sigma_{\mathrm{FAMILY}_{i}} \right)$$

$$\sigma_{\mathrm{AREA}_{i}}, \sigma_{GRID.\mathrm{ID}_{i},}\sigma_{\mathrm{FAMILY}_{i}}, \sigma_{\mathrm{spline}}\sim N\left( 0, 2 \right)$$

For each language *i*, we estimated the probability of the observed feature state across multiple categories. $\alpha$ is again the global baseline across all grid cells. $\alpha_{j}$ indicates category-specific intercepts for each possible state of the feature. The terms $\alpha_{\mathrm{AREA}_{i,j}}$, $\alpha_{GRID.\mathrm{ID}_{i,j}}$ and $\alpha_{\mathrm{FAMILY}_{i, j}}$ allow the intercept for each category to vary by area, grid cell, and family, respectively. The term ${t2}_{j}(\mathrm{lon}_{GRID.\mathrm{ID}_{i}},\mathrm{lat}_{GRID.\mathrm{ID}_{i}})$ represents a tensor product spline smooth for each category that captures geographic variation over the location of grid cell of the language (coordinates in Equal Earth Asia-Pacific projection EPSG:8859). All category-specific intercepts, varying effects, and tensor smooth term standard deviations followed normal priors with mean 0 and standard deviation 2 for each category.

We then extracted draws from the expected value of the posterior predictive distribution of each of these models to compute feature-wise estimates of the normalized Shannon entropy (H_n_) of the probabilities of all feature states per grid cell:

H_n_ = $-\frac{\sum_{j=1}^{J} \pi_{j}log(\pi_{j})}{log(J)}$

We collected the posterior means and standard deviations of H_n_ and of logit(H_n_) per feature and grid cell, separately for each dataset (TLI, GBI) and grid resolution. The models described in this section therefore act as “parametric bootstraps” to provide regularized estimates (with uncertainty) of grid-level entropy.

Predictor data aggregation

Predictor data (for an overview, see table S1) for our main models (see below) was aggregated at the level of the grid cells following an existing pipeline (*17*). For the (log) language richness predictor, we used the cell-wise logarithm of the count of languages and dialects according to Glottolog, v. 5.0 (*65*). For the (log) taxonomic diversity predictor, we used the cell-wise logarithm of the taxonomic diversity of the locally attested languages and dialects according to the Glottolog taxonomy (*65*) The taxonomic diversity index used is based on the index adopted in the R package densify (*66*). It adjusts for uneven taxonomic depths and node occurrences, producing a single diversity score based on balanced counts across taxonomy levels, with more diverse and balanced structures yielding higher scores. As environmental predictors, we used the centered and scaled first and second principal components of a probabilistic principal component analysis (PPCA) over eleven environmental variables from various sources (table S1) (*17*, *67*, *68*), performed in R using the pca() function of the pcaMethods package (*69*). Variables were extracted for the year 2000 CE, or, for the variables from WorldClim2 (*67*), for the averages for the years 1970–2000. The first PC of the environmental predictors had an R^2^ of 34.8 at the main spatial resolution and an R^2^ value of 35.2 at the resolution of the sensitivity analysis. Its main loadings were the number of months with mean temperature > 15°C (27.3% at both resolutions), mean annual temperature (23.7% in the main analysis, 23.4% in the sensitivity analysis) and temperature of the warmest quarter (22.7% at both resolutions). The second PC had an R^2^ of 18.6 in the main analysis and an R^2^ of 17.1 in the sensitivity analysis. Its main loadings were altitude (26.6% and 27.6% for the main and sensitivity resolutions, respectively), altitude variation (20.1% and 19.3%, respectively) and seasonal variance of precipitation (15.6% and 16.8%, respectively). For the (log) population density predictor, we used the cell-wise logarithm of the median population density in the year 2000 CE extracted from HYDE, v.3.3 (*68*). For the genetic predictor, we used the cell-wise median coefficient *F* (also known as Wright’s *F*) of 5737 unrelated individuals from 650 genetic populations with a sampling location that is sufficiently clear (median: 8 individuals per population). These populations represent 446 languages, as drawn from the GeLaTo (‘Genes and Languages Together’) database (*1*). *F* was calculated with PLINK v 1.9 (*70*) for each individual in the dataset as the observed (*H*_O_) minus expected (*H*_E_) homozygous genotypes by the number of non-missing genotypes (i.e., the number of non-missing SNPs) minus the expected number of homozygotes:

$$F=\frac{H_{O}-H_{E}}{N_{\mathrm{NM}}-H_{E}}$$

Finally, random effects by area were defined according to language assignments (Graff et al., in press) to one of the ten continent-sized areas (Africa, W and SW Eurasia, N-C Asia, S/SE Asia, New Guinea and Oceania, Australia, W N America, E N America, C America and S America) delineated in the AUTOTYP database (*31*).

Modelling structural diversity

As the response variable of our main models, again implemented in the brms (*61*) interface to Stan (*62*, *63*), we used the logit-transformed posterior mean entropies per grid cell ${(logit(H_{n}}_{i}))$ and the corresponding standard deviation of these estimates ${(sd(logit(H_{n}}_{i}))$ from the features of TLI (N=333) and GBI (N=196) separately. In our models, the total variance of each observation was composed of the observed standard deviation and an additional residual standard deviation $\sigma,$ which was estimated by the model, following a normal prior with mean 0 and standard deviation 1.

$${logit(H_{n}}_{i})\sim N\left( \eta_{i}, \sigma_{i} \right)$$

$$\sigma_{i}=\sqrt{\mathrm{sd}\left( {logit(H_{n}}_{i}) \right)^{2}+\sigma^{2}}$$

$$\sigma=N(0,1)$$

Our null model m1, including only (log) language richness and (log) taxonomic diversity as main predictors, was defined as follows:

$$\eta_{i}=\alpha+\alpha_{\mathrm{FEATURE}_{i}}+\alpha_{\mathrm{AREA}_{i}}+\alpha_{GRID.\mathrm{ID}_{i}}+$$

$${(\beta_{R}+\beta}_{R,\mathrm{FEATURE}_{i}}+\beta_{R,\mathrm{AREA}_{i}})\times R_{i}+$$

$$\left( {\beta_{T}+\beta}_{T,\mathrm{FEATURE}_{i}}+\beta_{T,\mathrm{AREA}_{i}} \right)\times T_{i}+$$

$$t2\left( \mathrm{lon}_{GRID.\mathrm{ID}_{i}},\mathrm{lat}_{GRID.\mathrm{ID}_{i}} \right)$$

$$\left( \begin{matrix} \alpha_{\mathrm{FEATURE}} \\ \beta_{R,FEATURE} \\ \beta_{T,FEATURE} \end{matrix} \right)\sim mvN\left( \left( \begin{matrix} 0 \\ 0 \\ 0 \end{matrix} \right),\Sigma_{\mathrm{FEATURE}} \right)$$

$$\Sigma_{\mathrm{FEATURE}}=\left( \begin{matrix} \sigma_{\alpha_{\mathrm{FEATURE}}}^{2} & \sigma_{\alpha_{\mathrm{FEATURE}}}\sigma_{\beta_{R,FEATURE}}\rho& \sigma_{\alpha_{\mathrm{FEATURE}}}\sigma_{\beta_{T,FEATURE}}\rho\\ \sigma_{\alpha_{\mathrm{FEATURE}}}\sigma_{\beta_{R,FEATURE}}\rho& \sigma_{\beta_{R,FEATURE}}^{2} & \sigma_{\beta_{R,FEATURE}}\sigma_{\beta_{T, FEATURE}}\rho\\ \sigma_{\alpha_{\mathrm{FEATURE}}}\sigma_{\beta_{T,FEATURE}}\rho& \sigma_{\beta_{R,FEATURE}}\sigma_{\beta_{T,FEATURE}}\rho& \sigma_{\beta_{T,FEATURE}}^{2} \end{matrix} \right)$$

$$\left( \begin{matrix} \alpha_{\mathrm{AREA}} \\ \beta_{R,AREA} \\ \beta_{T,AREA} \end{matrix} \right)\sim mvN\left( \left( \begin{matrix} 0 \\ 0 \\ 0 \end{matrix} \right),\Sigma_{\mathrm{AREA}} \right)$$

$$\Sigma_{\mathrm{AREA}}=\left( \begin{matrix} \sigma_{\alpha_{\mathrm{AREA}}}^{2} & \sigma_{\alpha_{\mathrm{AREA}}}\sigma_{\beta_{R,AREA}}\rho& \sigma_{\alpha_{\mathrm{AREA}}}\sigma_{\beta_{T,AREA}}\rho\\ \sigma_{\alpha_{\mathrm{AREA}}}\sigma_{\beta_{R,AREA}}\rho& \sigma_{\beta_{R, AREA}}^{2} & \sigma_{\beta_{R,AREA}}\sigma_{\beta_{T,AREA}}\rho\\ \sigma_{\alpha_{\mathrm{AREA}}}\sigma_{\beta_{T,AREA}}\rho& \sigma_{\beta_{R,AREA}}\sigma_{\beta_{T,AREA}}\rho& \sigma_{\beta_{T,AREA}}^{2} \end{matrix} \right)$$

$\alpha$ indicates the intercept, β_R_ indicates the fixed effect of the centered and scaled logarithm of language richness (R). β_T_ indicates the fixed effect of the centered and scaled logarithm of taxonomic diversity (T). The terms $\alpha_{\mathrm{FEATURE}_{i}}$, $\alpha_{\mathrm{AREA}_{i}}$ and $\alpha_{GRID.\mathrm{ID}_{i}}$ indicate that the intercept varies by feature, by the area of the grid cell and by grid cell. $\beta_{R,\mathrm{FEATURE}_{i}}$ and $\beta_{T,\mathrm{FEATURE}_{i}}$ indicate that both the richness and the taxonomic diversity slopes vary by feature. $\beta_{R,\mathrm{AREA}_{i}}$ and $\beta_{T,\mathrm{AREA}_{i}}$ indicate that the richness and the taxonomic diversity slopes further vary by area$.$ Lastly, the term $t2\left( \mathrm{lon}_{GRID.\mathrm{ID}_{i}},\mathrm{lat}_{GRID.\mathrm{ID}_{i}} \right)$ indicates a tensor product spline for smooths over the geographic location of the grid cell (coordinates in Equal Earth Asia-Pacific projection EPSG:8859).

We set the following priors on model parameters:

$$\alpha{, \beta}_{R},\beta_{T}\sim N\left( 0, 1 \right)$$

$$\alpha_{\mathrm{FEATURE}_{i}},\beta_{R,\mathrm{FEATURE}_{i}},\beta_{T,\mathrm{FEATURE}_{i}}\sim N\left( 0, \sigma_{\mathrm{FEATURE}_{i}} \right)$$

$$\alpha_{\mathrm{AREA}_{i}},\beta_{R,\mathrm{AREA}_{i}},\beta_{T,\mathrm{AREA}_{i}}\sim N\left( 0, \sigma_{\mathrm{AREA}_{i}} \right)$$

$$\alpha_{GRID.\mathrm{ID}_{i}}\sim N\left( 0, \sigma_{GRID.\mathrm{ID}_{i}} \right)$$

$$\sigma_{\mathrm{FEATURE}_{i}}, \sigma_{\mathrm{AREA}_{i},}\sigma_{GRID.\mathrm{ID}_{i}}, \sigma_{\mathrm{spline}}\sim N\left( 0,1 \right)$$

$$\mathbf{R}_{\mathbf{FEATURE}},\mathbf{R}_{\mathbf{AREA}}\sim LKJ\left( 2 \right)$$

The intercept and each of the fixed effects were assumed to follow a normal prior with mean 0 and standard deviation 1. Each varying effect term also individually followed a normal prior with mean 0 and standard deviation 1. Jointly, the varying effects by feature followed multivariate normal priors with mean 0 and variance-covariance $\Sigma_{\mathrm{FEATURE}}$. The effects by area followed multivariate normal priors with mean 0 and variance-covariance $\Sigma_{\mathrm{AREA}}$. The co-variance matrices were decomposed into a prior standard deviation vector and a correlation matrix **R**. Each **R** matrix was individually drawn from an *LKJ(*2) prior. The tensor smooth term standard deviation followed a normal prior with mean 0 and standard deviation 1.

Models m2-m8 followed the same structure of model m1. In addition to all terms in m1, their linear predictors additionally included different combinations of further linear main effects (P1, P2, D and *F*) and corresponding varying slopes by feature ($\beta_{P1,\mathrm{FEATURE}_{i}},\beta_{P2,\mathrm{FEATURE}_{i}},\beta_{D,\mathrm{FEATURE}_{i}},\beta_{F,\mathrm{FEATURE}_{i}}$) and area ($\beta_{P1,\mathrm{AREA}_{i}}, \beta_{P2,\mathrm{AREA}_{i}},\beta_{D,\mathrm{AREA}_{i}},\beta_{F,\mathrm{AREA}_{i}}$ (table S2).

Prior configurations in models m2-m8 were equivalent to m1, with sigma, the intercept, each fixed effect term, each varying effect term and the tensor smooth term standard deviation individually following a normal prior with mean 0 and standard deviation 1. Effects by feature and area each followed multivariate normal priors with mean 0 and variance-covariance $\Sigma_{\mathrm{FEATURE}}$and $\Sigma_{\mathrm{AREA}}$ respectively. These variance-covariance matrices followed the structure of $\Sigma_{\mathrm{FEATURE}}$and $\Sigma_{\mathrm{AREA}}$ from m1, but were appropriately re-dimensioned to include standard deviation vectors and correlation matrices **R** accounting for all varying effects included.

In addition to all terms in m1, m2 included all terms relating to the environment (i.e., P1 and P2 and all related terms). m3 additionally included all terms relating to (log) population density (i.e., D and all related terms). m4 additionally included all terms relating to individual genetic diversity (i.e., *F* and all related terms). m5 additionally included all terms relating to the environment and population density (i.e., P1, P2 and D, and all related terms). m6 additionally included all terms relating to the environment and genetics (i.e., P1, P2 and *F*, and all related terms). m7 additionally included all terms relating to population density and genetics (i.e., D and *F*, and all related terms). Finally, m8, the full model, additionally included all terms relating to the environment, population density and genetics (i.e., P1, P2, D, *F,* and all related terms). Table 1 provides an overview of models m1-m8 in R notation.

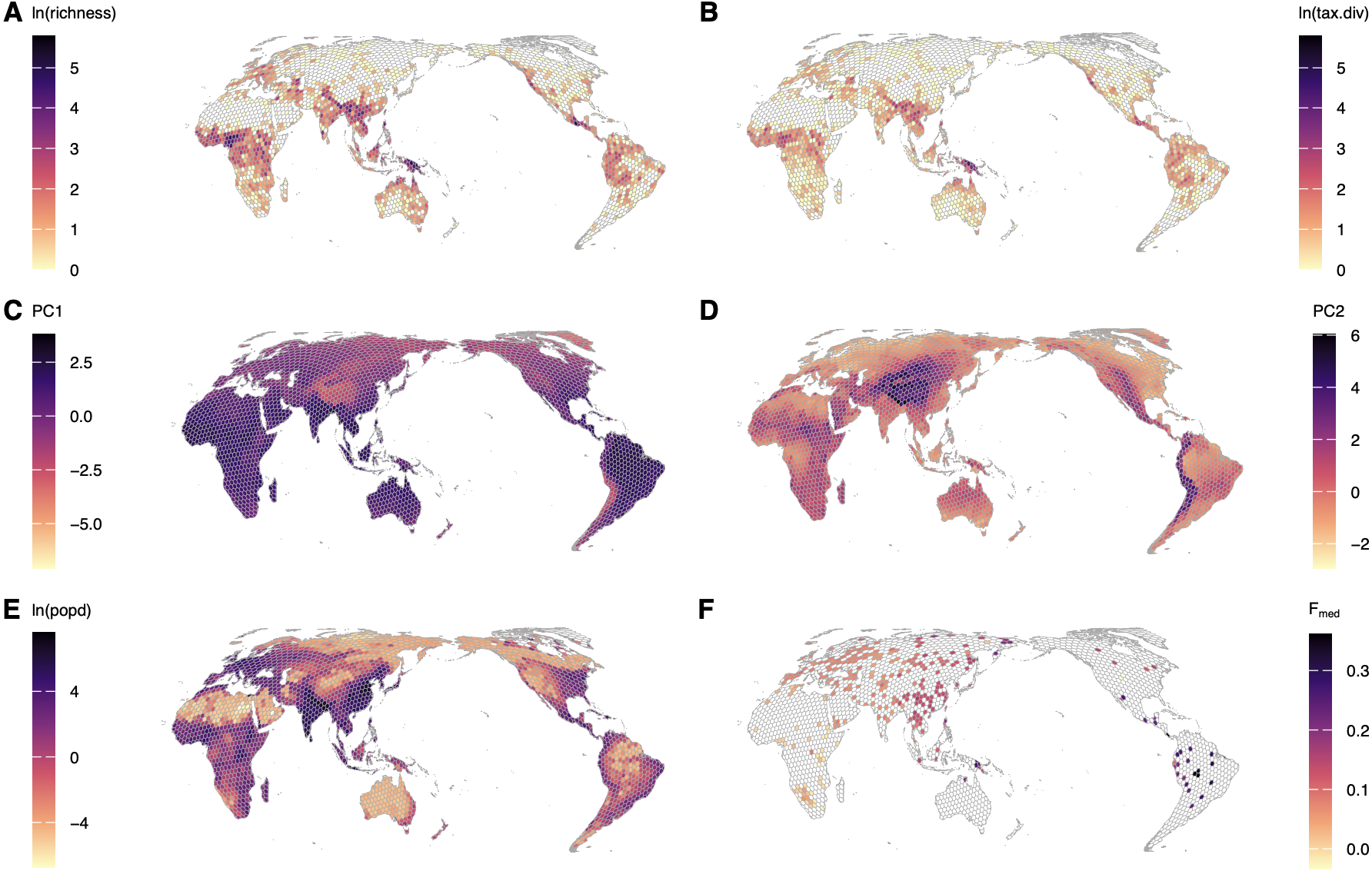

fig. S1.

Environmental and demographic predictors used for the sensitivity analyses models of local structural diversity at grid cell diameter 300 km. (**A**) (Log) language richness, i.e. number of languages and dialects per cell. (**B**) (Log) taxonomic diversity of languages and dialects per cell, scoring how diverse and balanced the included languages’ taxonomic relationships are. (**C**) First PC of environmental predictors (R2 = 35.2). Main loadings are the number of months with mean temperature > 15°C (27.3%), mean annual temperature (23.4%) and temperature of the warmest quarter (22.7%). (**D**) Second PC of environmental predictors (R2 = 17.1). Main loadings are altitude (27.6%), altitude variation (19.3%) and seasonal variance of precipitation (16.8%). (**E**) (Log) population density. (**F**) Median Wright’s *F* coefficient, with increasing values indicating higher excess homozygosity and therefore lower levels of genetic diversity.

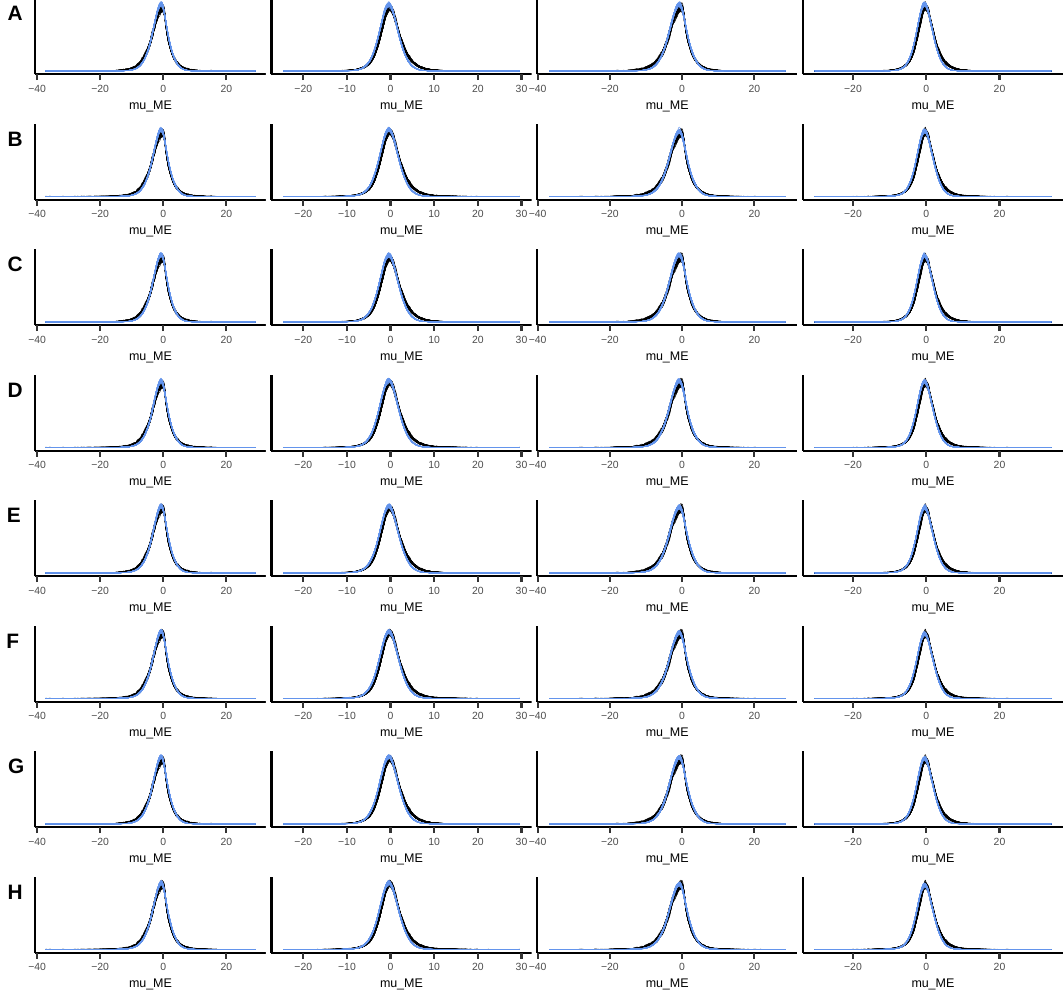

fig. S2.

Posterior predictive checks of all main analysis models, with 100 draws from feature-wise entropy predictions (feature models) in black and 100 predicted values including measurement error in blue. From left to right: TLI dataset, grid cell diameter 500 km; GBI dataset, grid cell diameter 500 km; TLI dataset, grid cell diameter 300 km; GBI dataset, grid cell diameter 300 km. (**A**) model m1. (**B**) model m2. (**C**) model m3. (**D**) model m4. (**E**) model m5. (**F**) model m6. (**G**) model m7. (**H**) model m8.

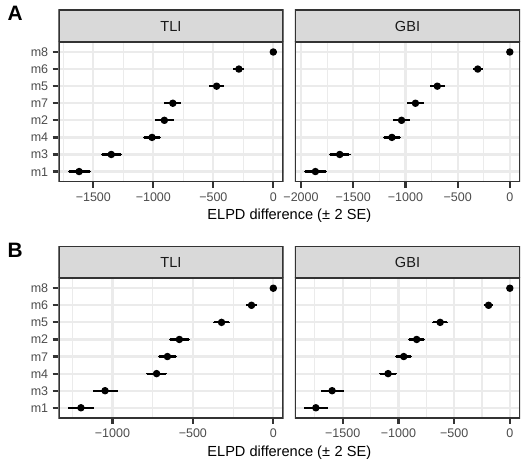

fig. S3.

Model comparison using ELPD differences between all main analysis models m1-m8 for the TLI-statistical and GBI-statistical linguistic datasets. (**A**) Grid cell diameter = 500 km. (**B**) Sensitivity analysis, grid cell diameter = 300 km. The full model m8 including environmental as well as (log) population density and genetic predictors performs best in both datasets and in both resolutions.

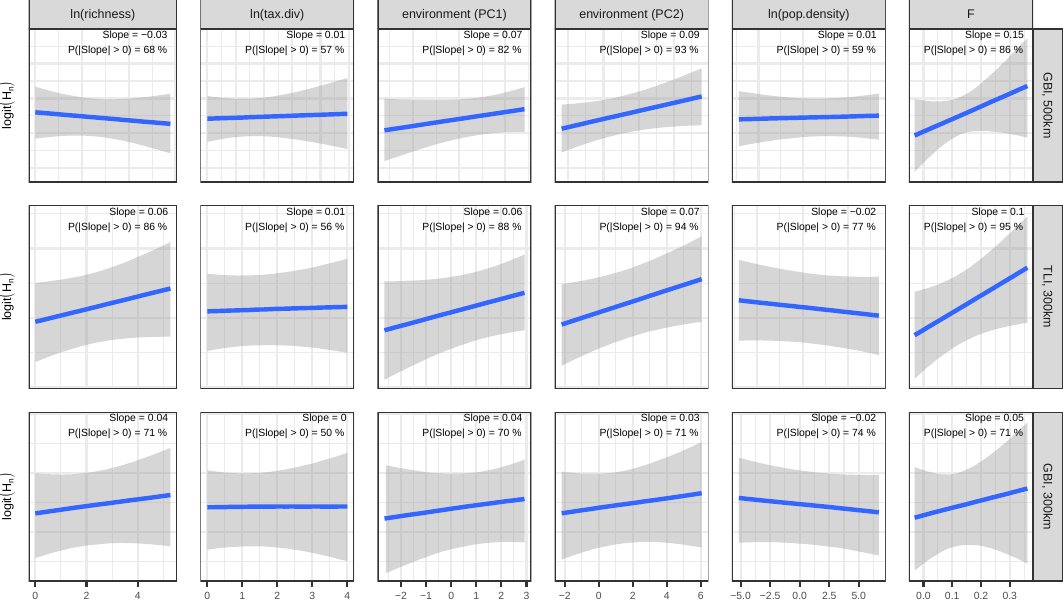
fig. S4.

Marginal effects at the mean of the six main predictors in the sensitivity analyses, i.e. with the GBI dataset at both geographic resolutions, and with the TLI dataset at the finer geographic resolution (grid cell 300 km). Intervals correspond to the 89% credible interval.

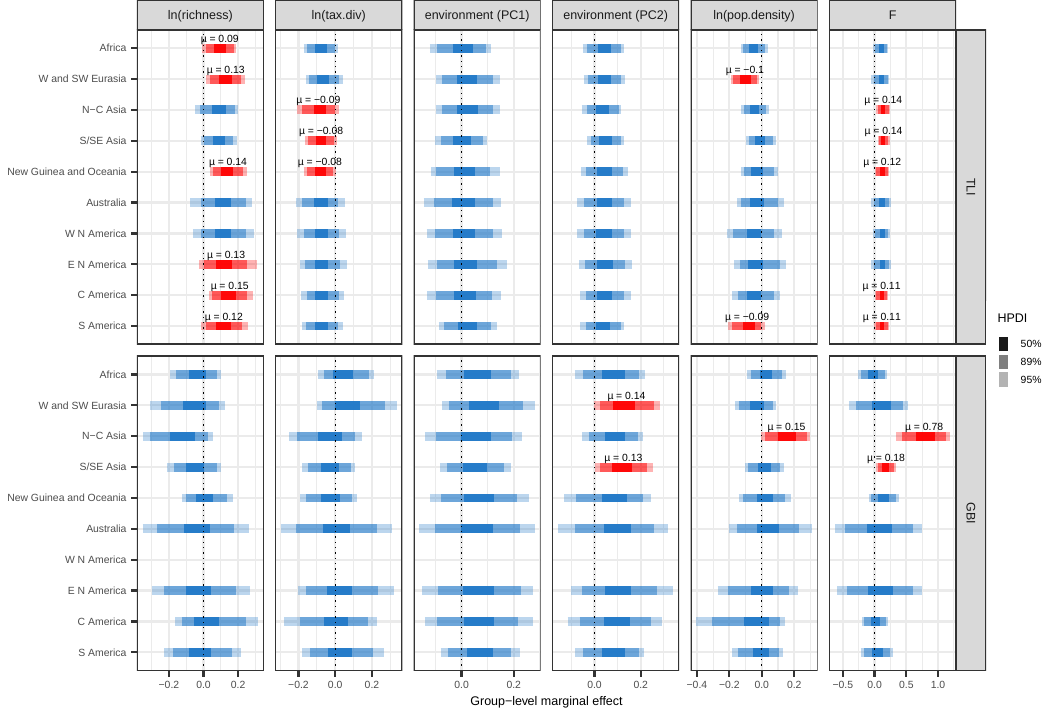
fig. S5.

Group-level marginal effects (slopes) of the six main predictors by geohistorical area as identified by AUTOTYP (*31*) in each dataset at the main analysis resolution (cell diameter = 500 km). Predictors with ≥ 95% posterior probability are colored red. For these predictors, µ reports the posterior mean slope.

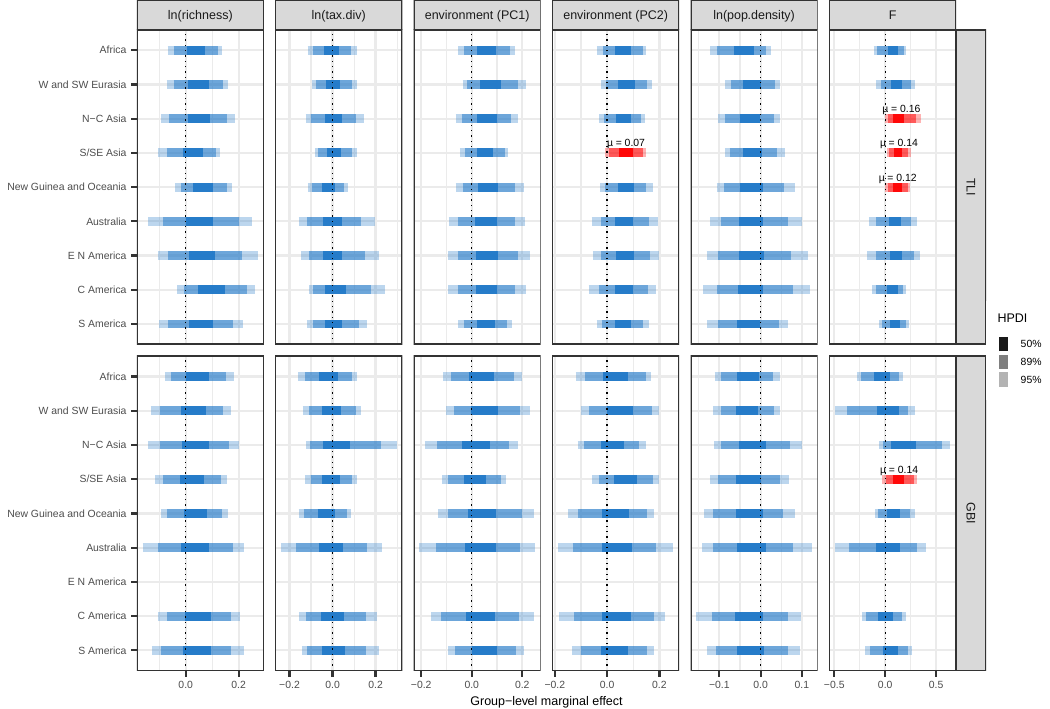
fig. S6.

Group-level marginal effects (slopes) of the six main predictors by geohistorical area as identified by AUTOTYP (*31*) in each dataset at the sensitivity analysis resolution (cell diameter = 300 km). Same plotting conventions as in fig. S5.

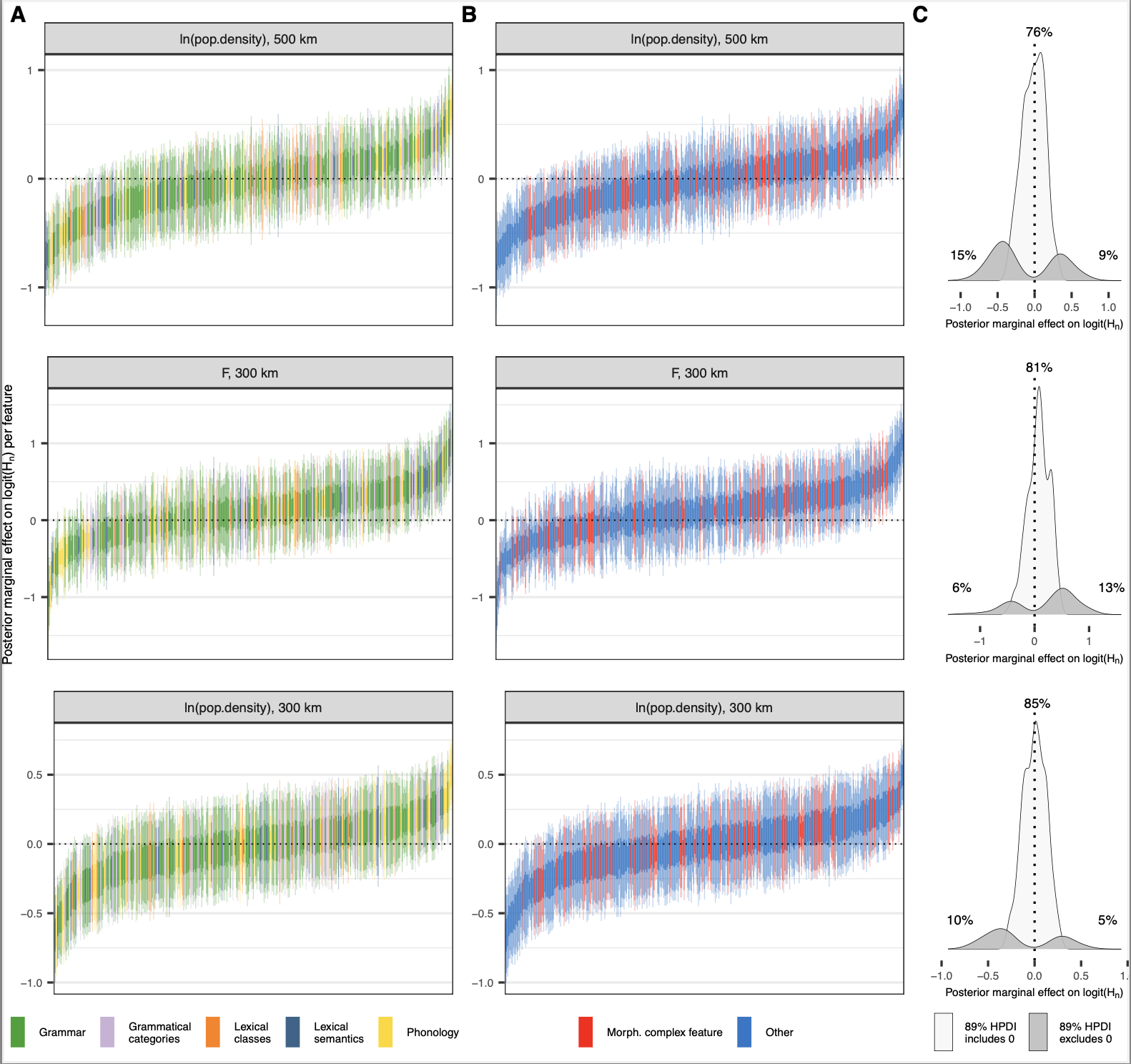
fig. S7.

Group-level marginal effects (slopes) by feature in GBI for *F* and log population density at both geographic resolutions. Same plotting conventions as in Figure 4. In (**B**) features on morphological complexity (in particular, the degree of fusion) (*30, 60*) are colored red. The list of features can be found in the OSF repository.

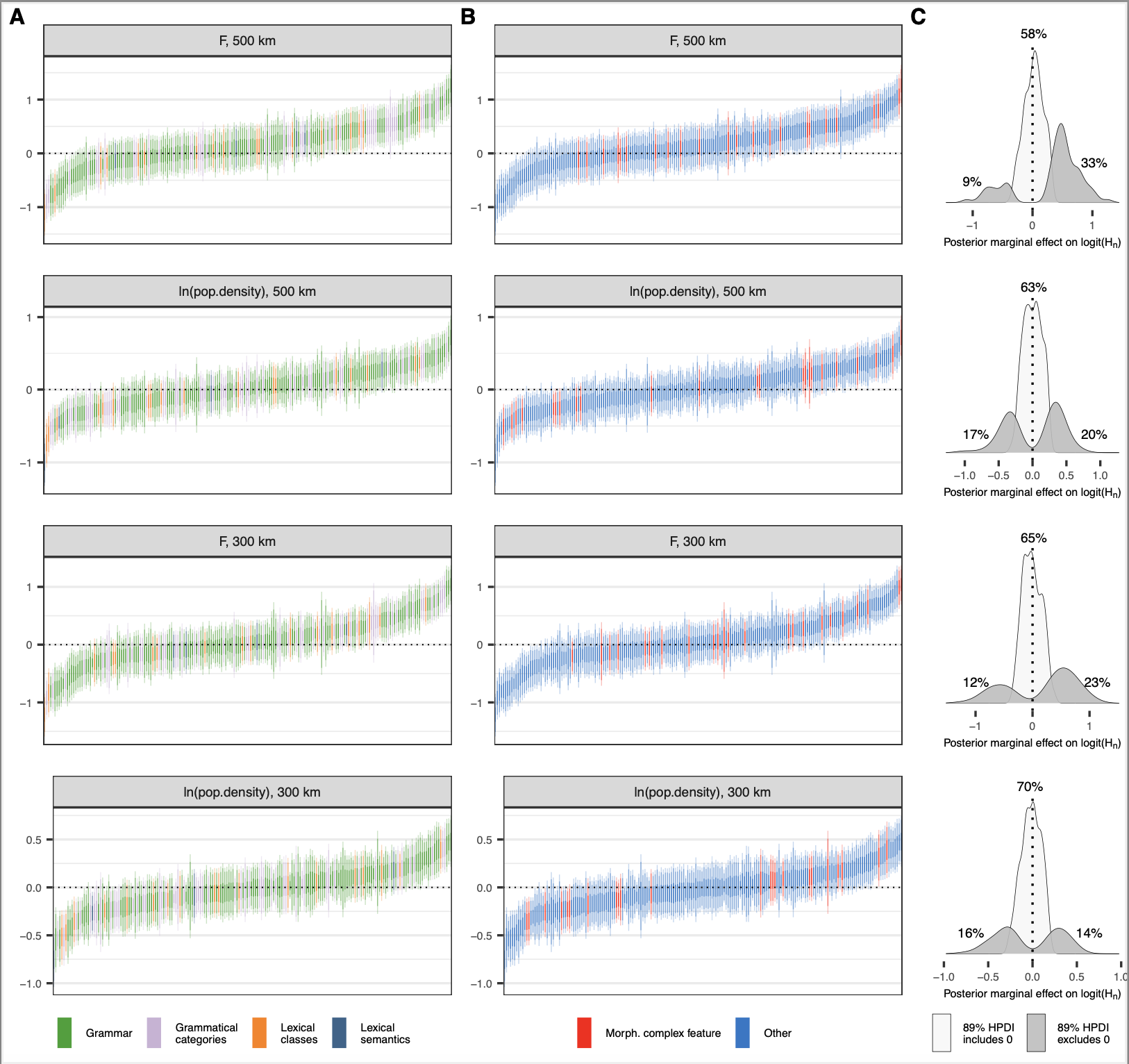
fig. S8.

Meta-analysis of contact effects on features across domains of language under different types of Group-level marginal effects (slopes) by feature in GBI for *F* and log population density at both geographic resolutions. Same plotting conventions as in Figure 4. In (**B**) features on morphological complexity (in particular, the degree of fusion) (*30, 60*) are colored red. The list of features can be found in the OSF repository.

table S1.

List of variables and their source.

| **#** | **Variable** | **Description*** | **Aggregation per cell** | **Source** |
| --- | --- | --- | --- | --- |
| 1 | ln.richness | Logarithm of language and dialect richness | log(count) | Glottolog, v. 5.0^61^ |
| 2 | ln.tax.div | Logarithm of taxonomic diversity of attested languages and dialects | see ‘Methods’ | Glottolog, v. 5.0^61^ |
| 3 | env.PC1 | First PC of a PCA over variables 11-21 (see ‘Methods’), main loadings relate to temperature | – | – |
| 4 | env.PC2 | Second PC of a PCA over variables 11-21 (see ‘Methods’), main loadings relate to terrain | – | – |
| 5 | ln.popd | Logarithm of population density (resolution: 5 arcminutes) | median | HYDE, v. 3.3^62^ |
| 6 | F.med | Wright’s *F* coefficient, excess of homozygosity | median | GeLaTo^1^ |
| 7 | feature | Linguistic feature identifier (features from TLI-statistical or GBI-statistical) | – | TLI/GBI^30^ |
| 8 | area | AUTOTYP area (10 “continent-sized” areas) associated with observation | – | TLI/GBI^30^ |
| 9 | grid.id | Identifier of a local observation unit | – | – |
| 10 | lon, lat | Coordinates in Equal Earth Asia-Pacific Projection (EPSG:8859) | – | – |
| 11 | precipitation | Seasonal variance of precipitation (resolution: 10 arcminutes). | median | WorldClim, v2^63^ |
| 12 | temp_mean | Mean annual temperature (resolution: 10 arcminutes). | median | WorldClim, v2^63^ |
| 13 | wettest | Precipitation in the wettest quarter (resolution: 10 arcminutes). | median | WorldClim, v2^63^ |
| 14 | warmest | Mean temperature of the warmest quarter (resolution: 10 arcminutes). | median | WorldClim, v2^63^ |
| 15 | n_warm_months | Months with mean temperature > 15°C (resolution: 10 arcminutes). | median | Derungs et al. 2018^17^ |
| 16 | grass | km2 of grassland/pasture (resolution: 5 arcminutes) | median | HYDE, v 3.3^62^ |
| 17 | crop | km2 of cropland (resolution: 5 arcminutes) | median | HYDE, v 3.3^62^ |
| 18 | elev | Highest elevation (resolution: 5 arcminutes) | max | Derungs et al. 2018^17^ |
| 19 | elevSD | Terrain ruggedness (sd of elevation, resolution: 5 arcminutes) | sd | Derungs et al. 2018^17^ |
| 20 | distOcean | Distance to ocean (resolution: 5 arcminutes) | median | Derungs et al. 2018^17^ |
| 21 | distRiver | Distance to river (resolution: 5 arcminutes) | median | Derungs et al. 2018^17^ |

* Variables from HYDE (3.3) were extracted for the year 2000 CE. Variables from WorldClim2 figure the averages for the years 1970-2000.

table S2.

Generalised additive mixed -effect models for estimating logit-transformed normalized entropy (div) using R notation.

| **Model** | **Description** | **Regression** |
| --- | --- | --- |
| m1 | Null model | bf(div \| se(div.sd, sigma = T)) ~ 1 + ln.richness + ln.tax.div +         (1 + ln.richness + ln.tax.div \| feature) +         (1 + ln.richness + ln.tax.div \| area) +         (1 \| grid.id) +         t2(lon, lat) ***** |
| m2 | Null model +  environment | m1 + env.PC1 + env.PC2 +        (env.PC1 + env.PC2 \| feature) +        (env.PC1 + env.PC2 \| area) |
| m3 | Null model +  population density | m1 + ln.popd +       (ln.popd \| feature) +        (ln.popd \| area) |
| m4 | Null model +  genetic diversity | m1 + F.med +        (F.med \| feature) +        (F.med \| area) |
| m5 | Null model +  environment +  population density | m1 + env.PC1 + env.PC2 + ln.popd +        (env.PC1 + env.PC2 + ln.popd \| feature) +        (env.PC1 + env.PC2 + ln.popd \| area) |
| m6 | Null model +  environment +  genetic diversity | m1 + env.PC1 + env.PC2 + F.med +        (env.PC1 + env.PC2 + F.med \| feature) +        (env.PC1 + env.PC2 + F.med \| area) |
| m7 | Null model +  population density +  genetic diversity | m1 + ln.popd + F.med +        (ln.popd + F.med \| feature) +        (ln.popd + F.med \| area) |
| m8 | Full model: null model +  environment+  population density +  genetic diversity | m1 + env.PC1 + env.PC2 + ln.popd + F.med +        (env.PC1 + env.PC2 + ln.popd + F.med \| feature) +        (env.PC1 + env.PC2 + ln.popd + F.med \| area) |

* The geographic locations of the grid cells (here: lon and lat) are indicated by their coordinates in the Equal Earth Asia-Pacific projection EPSG:8859.
